# Transcranial random noise stimulation of the primary visual cortex but not retina modulates visual contrast sensitivity

**DOI:** 10.1101/2022.02.28.482316

**Authors:** Weronika Potok, Alain Post, Marc Bächinger, Daniel Kiper, Nicole Wenderoth

## Abstract

Transcranial random noise stimulation (tRNS) has been shown to significantly improve visual perception. Previous studies demonstrated that tRNS delivered over cortical areas acutely enhances visual contrast detection of stimuli when tRNS intensity is optimized for the individual. However, it is currently unknown whether tRNS-induced signal enhancement could be achieved within different neural substrates along the retino-cortical pathway and whether the beneficial effect of optimal tRNS intensities can be reproduced across sessions.

In 3 experimental sessions, we tested whether tRNS applied to the primary visual cortex (V1) and/or to the retina improves visual contrast detection. We first measured visual contrast detection threshold (VCT; N=24, 16 females) during tRNS delivery separately over V1 (no tRNS, 0.75, 1, 1.5mA) and over the retina (no tRNS, 0.1, 0.2, 0.3mA), determined the optimal tRNS intensities for each individual (ind-tRNS), and retested the effects of ind-tRNS within the sessions. We further investigated whether we could reproduce the ind-tRNS-induced modulation on a different session (N=19, 14 females). Finally, we tested whether the simultaneous application of ind-tRNS to the retina and V1 causes additive effects.

We found that at the group level tRNS of 0.75mA decreases VCT compared to baseline when delivered to the V1. Beneficial effects of ind-tRNS could be replicated when retested within the same experimental session but not when retested in a separate session. Applying tRNS to the retina did not cause a systematic reduction of VCT, irrespective of whether the individually optimized intensity was considered or not. We also did not observe consistent additive effects of V1 and retina stimulation.

Our findings demonstrate that V1 seems to be more sensitive than the retina to tRNS-induced modulation of visual contrast processing.

**Significance statement:** Our findings confirm previous evidence showing acute online benefits of tRNS of V1 on visual contrast detection in accordance with the stochastic resonance phenomenon. We further extend it, demonstrating that the optimal tRNS intensity varies among participants, but when individually tailored it can improve visual processing when re-tested within the experimental session. The tRNS-induced enhancement in visual sensitivity seems to be specific for cortical contrast processing as stimulation of the retina did not lead to systematic effects.

## 1. Introduction

Transcranial random noise stimulation (tRNS) has been shown to significantly improve visual perception (see Potok et al., 2022 for review) when applied to visual cortex. Such performance improvements can manifest as both after-effects of visual training combined with tRNS (Contemori et al., 2019; Fertonani et al., 2011; Herpich et al., 2019; Pirulli et al., 2013) or acute effects during tRNS (Battaglini et al., 2020, 2019; Ghin et al., 2018; Pavan et al., 2019; van der Groen et al., 2019, 2018; van der Groen and Wenderoth, 2016). Studies exploring the acute effects of tRNS on visual processing have shown that noise stimulation of the primary visual cortex (V1) improves stimulus contrast detection, particularly, when visual stimuli are presented with near-threshold intensity (Battaglini et al., 2019; van der Groen and Wenderoth, 2016). What remains unknown is whether tRNS-induced signal enhancement, and related contrast sensitivity benefits, could be achieved at the retinal level. Modelling studies suggest noise benefits in retinal ganglion cells (Patel and Kosko, 2005) induced by both visual (Ghosh et al., 2009) and electrical noise (Wu et al., 2017). Moreover, previous research has suggested that the retina is susceptible to 8-20Hz alternating currents (Kar and Krekelberg, 2012; Schutter and Hortensius, 2010) which induce phosphenes even if the stimulation electrodes are placed over distal locations of the scalp (Laakso and Hirata, 2013; see Schutter, 2016 for review). Interestingly, improvement in vision was reported after repetitive transorbital alternating current stimulation at 5-30 Hz over the retina of patients with optic neuropathy or after optic nerve lesions (Fedorov et al., 2011; Gall et al., 2011, 2010; Sabel et al., 2011). They suggested that observed improvements were mediated by increased neuronal synchronization of residual structures and higher cortical areas within the visual system (Sabel et al., 2011). The retina and the optic nerve are interesting targets because they can be reliably reached even with low transcranial electrical stimulation intensities since the eyeball is an excellent conductor (Haberbosch et al., 2019). However, it remains unknown whether noise benefits resulting from tRNS can be induced at different levels of the retino-cortical processing pathway.

In this preregistered study, we investigated the effects of tRNS stimulation of the retina, primary visual cortex (V1) or both on visual detection performance.

## 2. Materials and methods

This study was preregistered on the Open Science Framework platform (https://osf.io/gacjw). The only difference to preregistered original plan concerns the included sample population. We stated that only participants who completed all three sessions will be included in our study.

During data collection not all the individuals who completed the 1^st^ and 2^nd^ sessions participated in the 3^rd^ session, due to the COVID-19 pandemic (Bikson et al., 2020). Nevertheless, we decided to keep all the data collected in session 1 and 2 (N=24) despite dropouts and lower sample size in session 3 (N=19, see *Participants* below).

### 2.1 Participants

Only individuals with no identified contraindications for participation according to established brain stimulation exclusion criteria (Rossi et al., 2009; Wassermann, 1998) were recruited for the study. All study participants provided written informed consent before the beginning of each experimental session. Upon study conclusion, they were debriefed and financially compensated for their time and effort. All research procedures were approved by the Cantonal Ethics Committee Zurich (BASEC Nr. 2018-01078) and performed in accordance with the Helsinki Declaration of the World Medical Association (2013 WMA Declaration of Helsinki).

The required sample size was estimated using an a priori power analysis (G*Power version 3.1; Faul, Erdfelder, Lang, & Buchner, 2007). Based on previous finding from van der Groen and Wenderoth (2016) we expected the effect of maximum contrast sensitivity improvement to correspond to Cohen’s d = 0.77. The power analysis revealed that fourteen participants should be included in an experiment to detect an effect of tRNS on contrast detection with repeated measures analysis of variance (rmANOVA, 4 levels of stimulation condition), alpha = 0.05, and 90% power, assuming the correlations among repeated measures = 0.5. However, there was no prior data available to investigate whether applying tRNS to two separate neural structures can cause additive effects. Therefore, we include more participants to ensure sufficient power. Moreover, this estimation hinges on the assumption that approx. 80% of the participants exhibit a behavioral response to tRNS (as indicated by Groen and Wenderoth, 2016). Thus, we collected data until N = 20 responders have been included. Responders were defined as individuals who exhibited improved detection in at least one tRNS condition in V1 and retina stimulation. Visual contrast detection is potentially prone to floor effects if the contrast detected at baseline approaches the technical limits of the setup. We decided to exclude participants that were exceptionally good in the visual task and present visual contrast threshold below 0.1 in the no tRNS baseline condition. We also excluded individuals with exceptional contrast threshold modulation (>100%) to avoid accidental results, e.g., due to participants responding without paying attention to the task. From the initially recruited sample of 32 participants, we excluded 8 individuals [5 participants had a contrast threshold below 0.1 in the baseline condition of one of the stimulation sessions (V1 or retina), 1 participant revealed exceptional contrast threshold modulation (>100%), 2 participants did not come back for the second session]. The final sample consisted of 24 healthy volunteers (16 females, 8 males; 24.4 ± 4.1, age range: 21-38) with normal or corrected-to-normal vision (see **Figure 1**A total number of 24 individuals participated in both 1^st^ (tRNS over V1) and 2^nd^ (tRNS over the retina) sessions. Due to the COVID-19 pandemic, we were forced to stop data collection for several months (Bikson et al., 2020). After returning to the lab, 5 participants dropped-out from the initial sample (2 had newly acquired contraindications for brain stimulation and 3 were not able to participate). 19 healthy volunteers (14 females, 5 males; 25.5 ± 5.2, age range: 21-39) were included into 3^rd^ session (tRNS over V1 and retina, see **Figure 1**).

**Figure 1.**
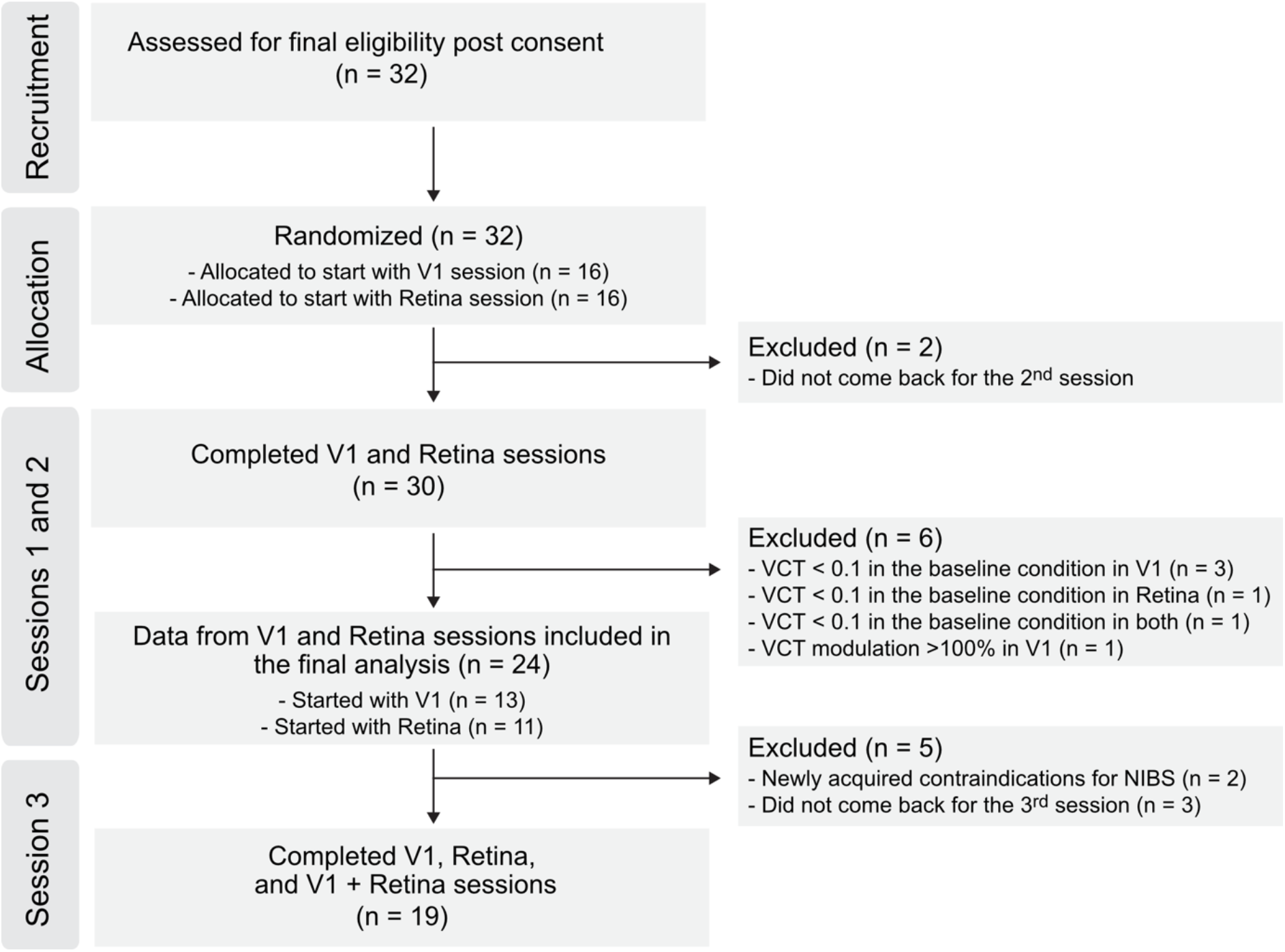
Flow diagram of the data collection progress through the phases of the study.

### 2.2 General Study design

To evaluate the influence of tRNS on visual contrast detection, we performed a series of three experimental sessions in which we delivered tRNS over different levels of the visual system, namely: V1, retina, or simultaneously over both V1 and retina (V1+Retina), during visual task performance (see **Figure 2A**). In each experiment, tRNS at low, medium and high intensity and a control no tRNS condition were interleaved in a random order (see *tRNS characteristics* below).

**Figure 2.**
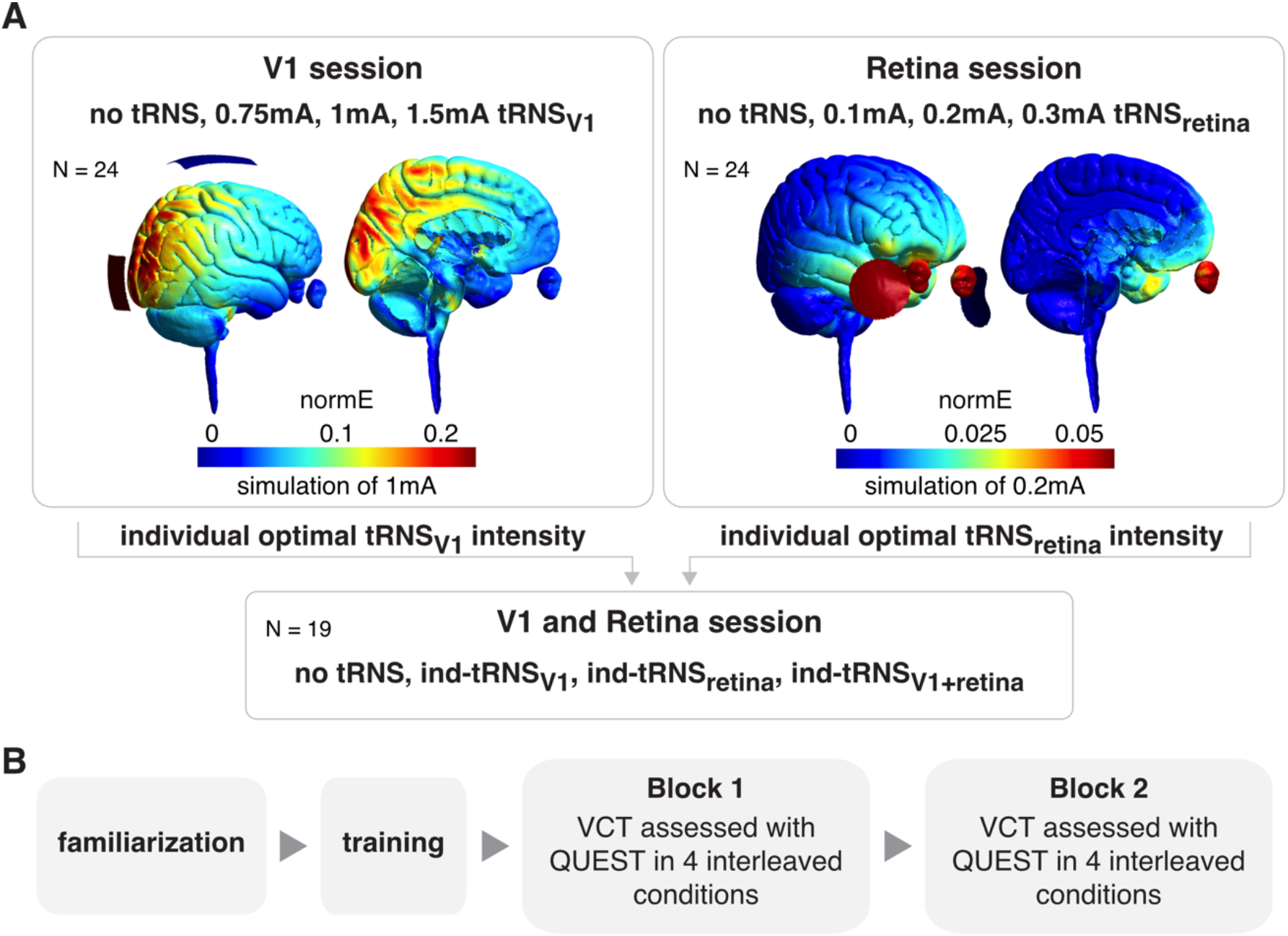
**A**. Experimental design and stimulation parameters. First, participants completed experimental sessions in which they received tRNS over V1 or retina (counterbalanced in order) in which the optimal individual tRNS intensity (ind-tRNS) was defined based on the behavioral performance. Next, the ind-tRNS was applied on the third session separately or simultaneously over V1 and retina. **B.** The order of measurements within each session. Each experimental session consisted of a familiarization protocol, followed by task training and two independent visual contrast threshold (VCT) assessments in 4 interleaved tRNS condition (as specified in A).

The order of experimental sessions for V1 and retina stimulation were counterbalanced across participants (13 participants started with V1 and 11 with retina stimulation). These experimental sessions took place on different days which were on average 2 weeks apart. Due to COVID-19 restrictions, the third session had to be delayed by 5 months on average.

Our main outcome parameter in all experimental sessions was a threshold of visual contrast detection (VCT) that was determined for each of the different tRNS conditions. VCT was independently estimated twice, in two separate blocks within each session (see **Figure 2B**). During the first two sessions we determined the individual optimal tRNS intensity (defined as the intensity causing the lowest VCT, i.e., biggest improvement in contrast sensitivity) for each participant in the V1 session (ind-tRNS_V1_) and the retina session (ind-tRNS_retina_). In the third session we then applied ind-tRNS_V1_ and ind-tRNS_retina_ to investigate the effect on VCT when V1 and retina are stimulated simultaneously.

#### 2.2.1 Visual stimuli

All experiments took place in a dark and quiet room, ensuring similar lighting conditions for all participants. Participants sat comfortably, 0.85m away from a screen, with their head supported by a chinrest. Visual stimuli were generated with Matlab (Matlab 2019b, MathWorks, Inc., Natick, USA) using the Psychophysics Toolbox extension (Brainard, 1997; Kleiner et al., 2007; Pelli, 1997) and displayed on a CRT computer screen (Sony CPD-G420). The screen was characterized by a resolution of 1280 x 1024 pixels, refresh rate of 85Hz, linearized contrast, and a luminance of 35 cd/m^2^ (measured with J17 LumaColor Photometer, Tektronix^™^). The target visual stimuli were presented on a uniform gray background in the form of a Gabor patch – a pattern of sinusoidal luminance grating displayed within a Gaussian envelope (full width at half maximum of 2.8 cm, i.e., 1° 53′ visual angle, with 7.3 cm, i.e., 4° 55′ presentation radius from the fixation cross). The Gabor patch pattern consisted of 16 cycles with one cycle made up of one white and one black bars (grating spatial frequency of 8 c/deg). Stimuli were oriented at 45° tilted to the left from the vertical axis (see **Figure 3B**), since it was shown that tRNS enhances detection of low contrast Gabor patches especially for non-vertical stimuli of high spatial frequency (Battaglini et al., 2020).

**Figure 3.**
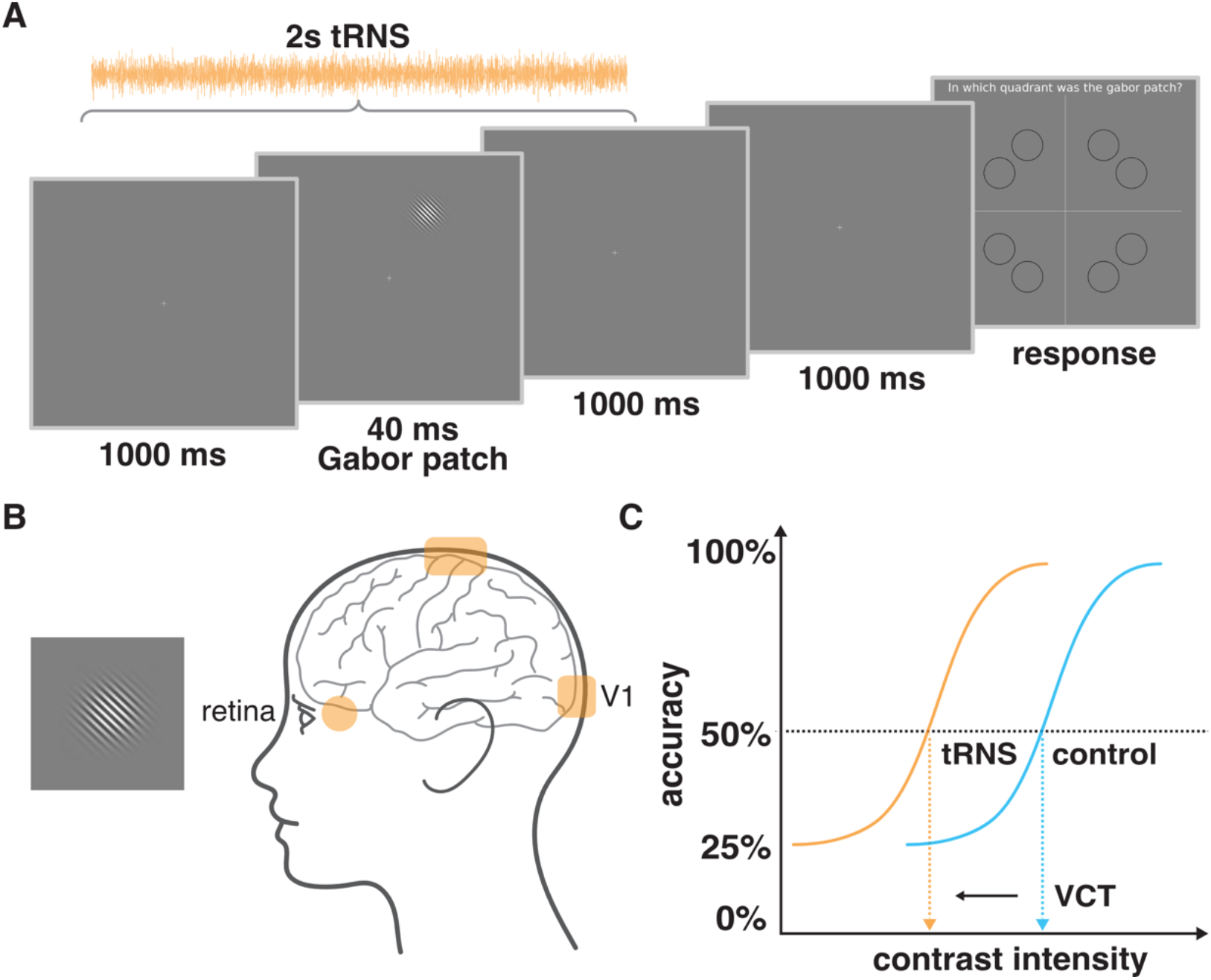
Experimental design. **A.** Example trial of 4-alternative forced choice task measuring visual contrast detection threshold (VCT). tRNS started 20ms after trial onset and was maintained for 2s **B.** Exemplary Gabor patch stimulus to be detected during the visual task and tRNS electrodes montage targeting V1 (rectangle) or retina (round, only the left side is shown but electrodes were mounted bilaterally). **C.** Example of dose-response psychometric curves and the detection of VCT for the 50% detection accuracy level. We hypothesize that the VCT will be lower (indicating better contrast detection performance of the participant) in one of the tRNS conditions (orange) than in the no tRNS control condition (blue).

#### 2.2.2 Four-alternative forced choice visual detection task

In all three experiments participants performed a four-alternative forced choice (4-AFC) visual task, designed to assess an individual VCT, separately for each tRNS condition. Such protocol was shown to be more efficient for threshold estimation than commonly used 2-AFC (Jäkel and Wichmann, 2006). In the middle of each 2.04s trial, a Gabor patch was presented for 40ms in one of the 8 locations (see **Figure 3A**). To account for potential differences in the extent to which tRNS affects different retinotopic coordinates and to avoid a spatial detection bias, the visual stimuli were presented pseudo-randomly and appeared the same number of times (20) in each of the eight locations on the screen within each experimental block (van der Groen and Wenderoth, 2016). The possible locations were set on noncardinal axes, as the detection performance for stimuli presented in this way is less affected (i.e. less variable) than when stimuli are positioned on the cardinal axes (Cameron et al., 2002; van der Groen and Wenderoth, 2016). The trial was followed by 1s presentation of fixation cross after which the ‘response screen’ appeared. Participants’ task was to decide in which quadrant of the screen the visual stimulus appeared and indicate its location on a keyboard. The timing of the response period was self-paced and not limited. Participants completed a short training (10 trials) at the beginning of each session, with the stimulus presented always at high contrast, in order to ensure that they understand the task (**Figure 2B**).

During the main experiment, VCT was estimated using the QUEST staircase maximum likelihood procedure (Watson and Pelli, 1983) implemented in the Psychophysics Toolbox in Matlab (Brainard, 1997; Kleiner et al., 2007; Pelli, 1997). The thresholding procedure starts with a presentation of the visual stimulus displayed with 0.5 contrast intensity (for visual contrast intensity range of minimum 0 and maximum 1). When participants answer correctly QUEST lowers the presented contrast intensity, when participants answer incorrectly QUEST increases the presented contrast. The estimated stimulus contrast is adjusted to yield 50% detection accuracy (i.e., detection threshold criterion, see **Figure 3C**). Note that for 4-AFC task 25% accuracy corresponds to a chance level. The remaining parameters used in the QUEST staircase procedure included: steepness of the psychometric function, beta = 3; fraction of trials on which the observer presses blindly, delta = 0.01; chance level of response, gamma = 0.25; step size of internal table grain = 0.001; intensity difference between the largest and smallest stimulus intensity, range = 1. VCT was assessed across 40 trials per tRNS condition (40 trials x 4 conditions x 2 blocks; total number of 320 trials per experimental session).

#### 2.2.3 tRNS characteristics

In tRNS trials, high-frequency tRNS (hf-tRNS, 100-640Hz) with no offset was delivered. The probability function of random current intensities followed a Gaussian distribution with 99% of the values lying between the peak-to-peak amplitude (Potok et al., 2022). Stimulation started 20ms after trial onset and was maintained for 2s (**Figure 3A**). Subsequently a fixation cross was displayed for 1 s, followed by the self-paced response time. tRNS waveforms were created within Matlab (Matlab 2020a, MathWorks, Inc., Natick, USA) and sent to a battery-driven electrical stimulator (DC-Stimulator PLUS, NeuroConn GmbH, Ilmenau, Germany), operated in REMOTE mode, via a National Instruments I/O device USB-6343 X series, National Instruments, USA). The active tRNS conditions and no tRNS control condition were interleaved and presented in random order. Timing of the stimuli presentation, remote control of the tRNS stimulator, and behavioral data recording were synchronized via Matlab (Matlab 2020a, MathWorks, Inc., Natick, USA) installed on a PC (HP EliteDesk 800 G1) running Windows (Windows 7, Microsoft, USA) as an operating system. The impedance between the electrodes was monitored and kept below 15 kΩ. For all experiments we used electric field modelling to ensure optimal electrode placement. All simulations were run in SimNIBS 2.1 (Thielscher et al., 2015) using the average MNI brain template (**Figure 2A**). The software enables finite-element modelling of electric field distribution of direct current stimulation without taking into account the temporal characteristics. Note, that the software enables simulation of electric field within the brain and eyeball but does not include the optic nerve.

Before the start of the main experiment, participants were familiarized with tRNS and we assessed the detectability of potential sensations (**Figure 2B**). The detection task consisted of 20 trials. Participants received either 5s tRNS (0.75, 1, and 1.5mA tRNS in V1 session; 0.1, 0.2, and 0.3mA tRNS in the retina session; or ind-tRNS_V1_, ind-tRNS_retina_, ind-tRNS_V1+retina_ in V1+Retina session) or no tRNS. Their task after each trial was to indicate on a keyboard whether they felt something underneath the tRNS electrodes. The determined detection accuracy (hit rates, HR) of the cutaneous sensation induced by tRNS served as a control to estimate whether transcutaneous effects of the stimulation might have confounded the experimental outcomes (Potok et al., 2021).

##### V1 session – testing the effect of no, low, medium or high intensity tRNS targeting V1 on visual detection performance

In the V1 session, we asked whether tRNS over V1 modulates VCT. To target V1 we used an electrode montage that was previously shown to be suitable for V1 stimulation (Herpich, 2019; van der Groen and Wenderoth, 2016). One tRNS 5×5cm rubber electrode was placed over the occipital region (3 cm above inion, Oz in the 10-20 EEG system) and one 5×7cm rubber electrode over the vertex (Cz in the 10-20 EEG system). Electroconductive gel was applied to the contact side of the rubber electrodes (NeuroConn GmbH, Ilmenau, Germany) to reduce skin impedance.

tRNS was delivered with 0.75mA (low), 1mA (medium), and 1.5mA (high) amplitude (peak-to-baseline), resulting in maximum current density of 60 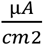, which is below the safety limits for transcranial electrical stimulation (Fertonani et al., 2015). tRNS power, corresponding to the variance of the electrical noise intensities distribution, was 0.109, 0.194 and 0.436mA^2^ in the 0.75, 1 and 1.5mA condition, respectively (Potok et al., 2022).

##### Retina session – testing the effect of no, low, medium or high intensity tRNS targeting the retina on visual detection performance

To further explore the influence of electrical random noise on visual processing we delivered tRNS over the retina during a visual contrast detection task. To stimulate the retina, face skin-friendly self-adhesive round electrodes with a diameter of 32mm (TENS-EMS pads Axion GmbH, Germany) were placed on the sphenoid bones of the right and left temples. Electroconductive gel was applied to the contact side of each electrode to additionally reduce skin impedance.

Dose-response effects were assessed with VCT during tRNS applied with the intensity of 0.1mA (low), 0.2mA (medium), and 0.3mA (high) amplitude (peak-to-baseline), resulting in a maximum current density of 29.3 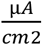, which is well below the safety limits for transcranial electrical stimulation (Fertonani et al., 2015). tRNS power, corresponding to the variance of the electrical noise intensities distribution, was 0.002, 0.008 and 0.017mA^2^ in the 0.1, 0.2 and 0.3mA condition, respectively (Potok et al., 2022). The selected intensities are commonly used in transorbital alternating current stimulation studies that have reported stimulation induced effects (Fedorov et al., 2011; Gall et al., 2011, 2010; Sabel et al., 2011). Furthermore, we had performed a pilot experiment (N = 30) to which assess a flickering threshold when low-frequency tRNS was used (0.1-100Hz). We found that flickering was perceived for a mean intensity of 0.146 ±0.08mA (peak-to-baseline) suggesting that the stimulation intensities chosen in this experiment should be suitable for reaching the retina. Our pilot experiment further revealed that flickering was induced by low-frequency tRNS but not by high-frequency tRNS (as used in the main experiments).

##### V1+Retina session – testing the additive effect of simultaneously applying tRNS to V1 and the retina on visual detection performance

The final experimental session aimed to investigate potential additive effects of delivering electrical random noise simultaneously to V1 and the retina on visual contrast sensitivity.

In this session, we combined the electrodes montages over V1 and the retina and applied tRNS with individual optimal intensities as determined in the first two experimental sessions (i.e., ind-tRNS_V1_ and ind-tRNS_retina_).

In the V1+Retina session, we compared the VCT in four conditions: (i) tRNS over V1 at its optimal intensity (ind-tRNS_V1_), (ii) tRNS over retina at its optimal intensity (ind-tRNS_retina_), (iii) simultaneous tRNS over V1 and the retina at their respective optimal intensities (ind-tRNS_V1+retina_), and (iv) no tRNS. All conditions were interleaved and presented in a randomized order.

### 2.3 Statistical Analysis

All the statistical analyses were preregistered and did not deviate from the original plan. Statistical analyses were performed in IBM SPSS Statistics version 26.0 (IBM Corp.). All data was tested for normal distribution using the Shapiro-Wilks test. Variance is reported as SD in the main text and as SE in the figures.

First, we tested whether baseline VCT in the no tRNS condition differed across the three experimental sessions using a Bayesian rmANOVA with the factor *time* (blocks 1-2 in sessions 1-3, i.e., six consecutive time points) using the Bayes factor testing for evaluation the absence versus presence of an effect.

For all rmANOVA models, sphericity was assessed with Mauchly’s sphericity test. The threshold for statistical significance was set at α = 0.05. Bonferroni correction for multiple comparisons was applied where appropriate (i.e., post hoc tests). Partial eta-squared (small 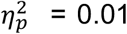, medium 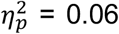, large 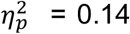; Lakens, 2013) values are reported as a measure of effect-sizes.

VCT data collected in the V1 session (tRNS_V1_) was analyzed with a rmANOVA with the factor *tRNS* (no, 0.75, 1, and 1.5mA tRNS) and the factor *block* (1^st^, 2^nd^). For each individual and each block, we determined the maximal behavioral improvement, i.e., lowest VCT measured when tRNS was applied, and the associated “optimal” tRNS intensity (ind-tRNS_V1_). The maximal behavioral improvements in the 1^st^ and the 2^nd^ block were compared using a t-test (2-tailed) for dependent measurements. We further tested whether ind-tRNS_V1_ of the 1^st^ and 2^nd^ block were correlated using Spearman’s rank correlation coefficient (because of categorical characteristics of ind-tRNS_V1_ variable). Importantly, we determined ind-tRNS_V1_ in the 1^st^ block, and then used the VCT data of the separate 2^nd^ block to test whether the associated VCT is lower compared to the no tRNS condition using t-tests for dependent measures. Since we had the directional hypothesis that VCT is lower for the optimal tRNS intensity compared to no tRNS this test was 1-tailed. Determining ind-tRNS_V1_ and testing its effect on VCT in two separate datasets is important to not overestimate the effect of tRNS on visual detection behavior.

VCT data collected in the Retina session (tRNS_retina_) was analyzed with a rmANOVA with the factor of *tRNS* (no, 0.1, 0.2, and 0.3mA tRNS) and the factor *block* (1^st^, 2^nd^). Again, for each individual and each block, we determined the maximal behavioral improvement and the associated ind-tRNS_retina_. We compared results obtained in the first and second block using the same statistical tests as for the V1 session. The maximal behavioral improvements were compared using a t-test (2-tailed) for dependent measurements. Correlation of ind-tRNS_retina_ of the 1^st^ and 2^nd^ block was tested using Spearman’s rank correlation coefficient. We examined whether the ind-tRNS_retina_ determined based on the best behavioral performance in 1^st^ block, caused VCT to be lower compared to the no tRNS condition when retested on the independent dataset (2^nd^ block) using t-tests (1-tailed) for dependent measures.

VCT data collected in the V1+Retina session (tRNS_V1+retina_) was analyzed with a rmANOVA with the factor *tRNS site* (ind-tRNS_V1_, ind-tRNS_retina_, ind-tRNS_V1+retina_, and no tRNS) and the factor *block* (1^st^, 2^nd^). Moreover, we compared behavioral improvement for ind-tRNS_V1_ and ind-tRNS_retina_ between sessions (tRNS_V1_ and tRNS_V1+retina_, tRNS_retina_ and tRNS_V1+retina_, respectively) using a Pearson correlation coefficient.

As a control analysis we repeated the main analyses of VCT (rmANOVA were we observed tRNS-induced significant difference) with adding cutaneous sensation as covariate (see *tRNS characteristics*).

## 3. Results

We first tested whether VCT measured during the no tRNS condition differed between the experimental sessions or blocks (i.e., six consecutive time points, see **Figure 4**). Bayesian rmANOVA with the factor *time* (1-6) revealed that the baseline VCT measured in the no tRNS condition did not differ over time (BF_10_ = 0.06, i.e., strong evidence for the H0) indicating that detection performance was rather stable across sessions.

**Figure 4.**
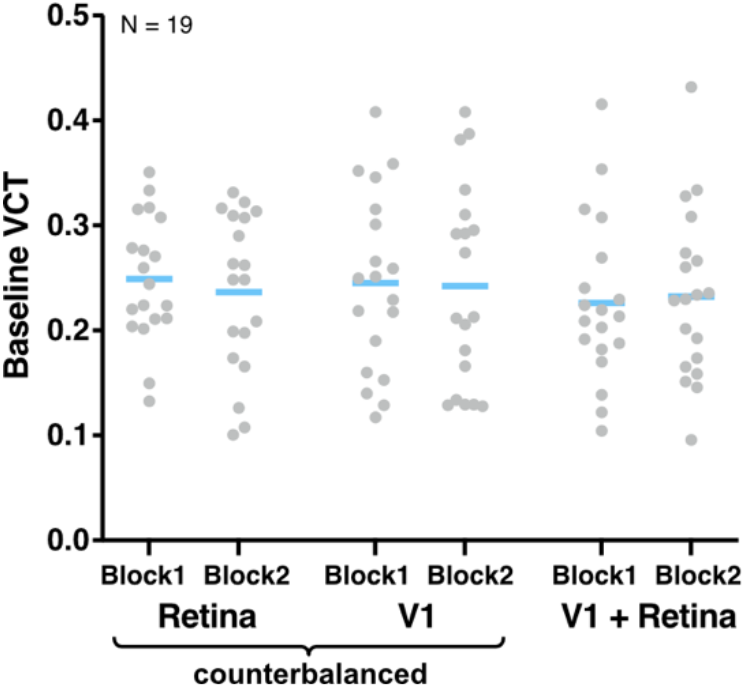
Baseline VCT measured in the no tRNS condition in both blocks in V1, Retina and V1+Retina sessions. Blue lines indicate mean, gray dots indicate single subject data.

### 3.1 tRNS over V1 modulates visual contrast threshold

In the V1 session, we investigated whether tRNS modulates the visual contrast detection when applied to V1. We measured VCT during tRNS_V1_ at intensities of 0.75, 1, to 1.5mA versus no tRNS control condition. We found a general decrease in VCT when tRNS was applied (*tRNS* main effect: F_(3, 69)_ = 4.54, p = 0.006, 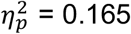) indicating that adding noise to V1 improved contrast sensitivity (**Figure 5A**). Post hoc comparisons revealed that the 0.75mA stimulation was most effective in boosting contrast processing, which differed significantly from the no tRNS control condition (p = 0.045, MD = −8.69 ±15.99%). As we observed that most of our participants could detect tRNS V1 conditions (mean HR = 76.04 ±22.16% measured via cutaneous sensations detection task) we reanalyzed our main outcome parameter by adding sensation detection HR as a covariate. The main effect of *tRNS* remained highly significant (F_(3, 66)_ = 4.17, p = 0.009, 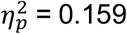), making it unlikely that cutaneous sensation was the main driver of our results. Neither the main effect of *block* (F_(1, 23)_ = 0.18, p = 0.678) nor *tRNS*block* interaction (F_(3, 69)_ = 0.82, p = 0.488) reached significance.

**Figure 5.**
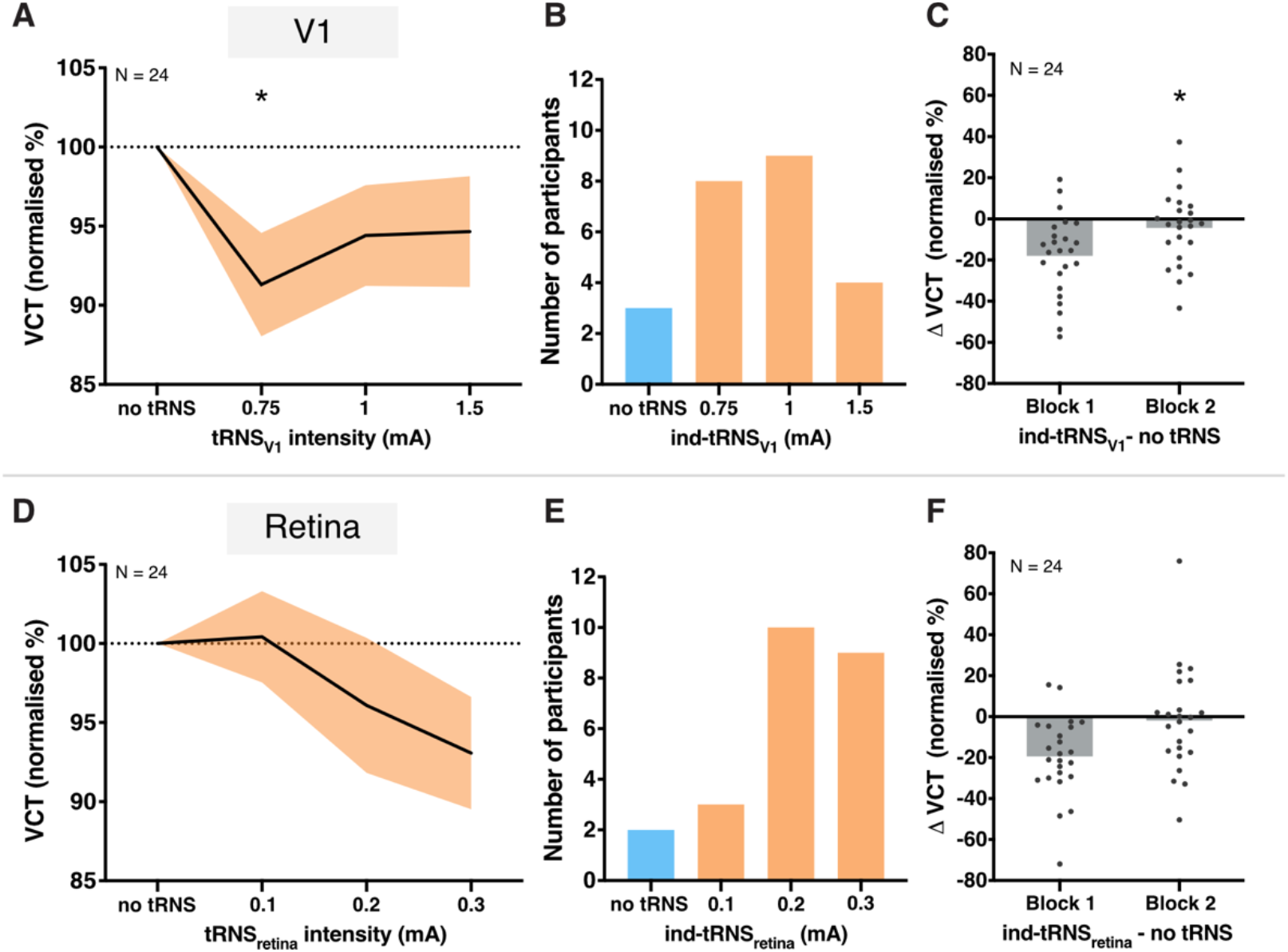
Results of V1 and Retina sessions. **A.** Effect of tRNS_V1_ on VCT on a group level measured across 1^st^ and 2^nd^ block in V1 session. Decrease in VCT reflects improvement of visual contrast sensitivity. All data mean ± SE. **B.** Individually defined optimal tRNS_V1_ based on behavioral performance during the 1^st^ block. **C.** Detection improvement effects of individualized tRNS_V1_. **D.** Effect of tRNS_retina_ on VCT on a group level measured across 1^st^ and 2^nd^ block in Retina session. All data mean ± SE. **E.**Individually defined optimal tRNS_retina_ based on behavioral performance during the 1^st^ block. **F.**Detection modulation during individualized tRNS_retina_. Gray dots indicate single subject data, gray bars indicate group mean; *p < 0.05.

When comparing tRNS-induced effects between the 1^st^ and 2^nd^ block we found that the maximal behavioral improvement (i.e. the maximal tRNS_V1_-induced lowering of the VCT relative to the no tRNS condition) differed only insignificantly between the 1^st^ (MD = −17.98 ±19.6%) and the 2^nd^ block (MD = −16.63 ±15.11%, t_(23)_ = −0.304, p = 0.764). However, participants’ optimal ind-tRNS_V1_ of block 1 and 2 (i.e., the tRNS intensity causing the largest VCT reduction in each block) were not correlated (rho = 0.225, p = 0.290).

Finally, we determined ind-tRNS_V1_ in the 1^st^ block (**Figure 5B**) and tested whether it caused a decrease in VCT compared to the no tRNS condition using the data of block 2. Indeed, VCT decreased in 15 out of 24 individuals (MD = −4.45 ±17.9%) and this effect reached statistical significance (t_(23)_ = 1.72, p = 0.049, **Figure 5C**). Note that the optimal ind-tRNS_V1_ intensity and the associated VCT effect were determined on independent data sets to avoid circularity.

### 3.2 tRNS over the retina does not modulate visual contrast threshold consistently

In the Retina session, we explored the effects of tRNS applied over the retina on visual contrast detection. VCT was measured during tRNS_retina_ at intensities of 0.1, 0.2, to 0.3mA versus no tRNS control condition. Even though, on the group level, we observed decrease in VCT with increasing tRNS_retina_ intensity (MD = −6.93 ±17.39% on average in the 1^st^ and 2^nd^ block for 0.3mA) the effect was not significant (F_(3, 69)_ = 1.69, p = 0.177, **Figure 5D**). There was also no main effect of *block* (F_(1, 23)_ = 0.04, p = 0.840) or *tRNS*block* interaction (F_(3, 69)_ = 0.55, p = 0.652). The maximal behavioral improvements, defined as the maximal tRNS_retina_-induced lowering of the VCT were not significantly different between the 1^st^ (MD = −19.44 ±19.43%) and the 2^nd^ (MD = −11.96 ±22.79%) block (t_(23)_ = −1.197, p = 0.243). Similar to the ind-tRNS_V1_, the optimal ind-tRNS_retina_ intensity defined in the 1^st^ and 2^nd^ block were not significantly correlated among participants (rho = 0.321, p = 0.126). The ind-tRNS_retina_ determined in the 1^st^ block (**Figure 5E**) did not significantly lower the VCT compared to the no tRNS condition when retested on the independent VCT dataset of block 2 (t_(23)_ = 1.05, p = 0.15, VCT decrease in 13 out of 24 individuals, MD = −1.89 ±25.29%).

### 3.3 No effects of simultaneous tRNS of V1 and retina on visual contrast threshold

The aim of V1+Retina session was to explore whether the effects of ind-tRNS_V1_ and ind-tRNS_retina_ determined in sessions 1 and 2 would have additive effects when combined during simultaneous V1 and retina stimulation (**Figure 6A**). Against our hypothesis, we did not observe a consistent decrease in VCT on the group level, neither when considering *tRNS site* (F_(3, 54)_ = 0.54, p = 0.660), *block* (F_(1, 18)_ = 2.73, p = 0.116) nor *tRNS site*block* interaction (F_(3, 54)_ = 0.31, p = 0.822). Although the simultaneous stimulation with ind-tRNS_V1+retina_ led to a decrease in VCT in the 1^st^ block (MD = −4.12 ±25.64%), this difference was not significant (t_(18)_ = 0.83, p = 0.21, **Figure 6B**).

**Figure 6.**
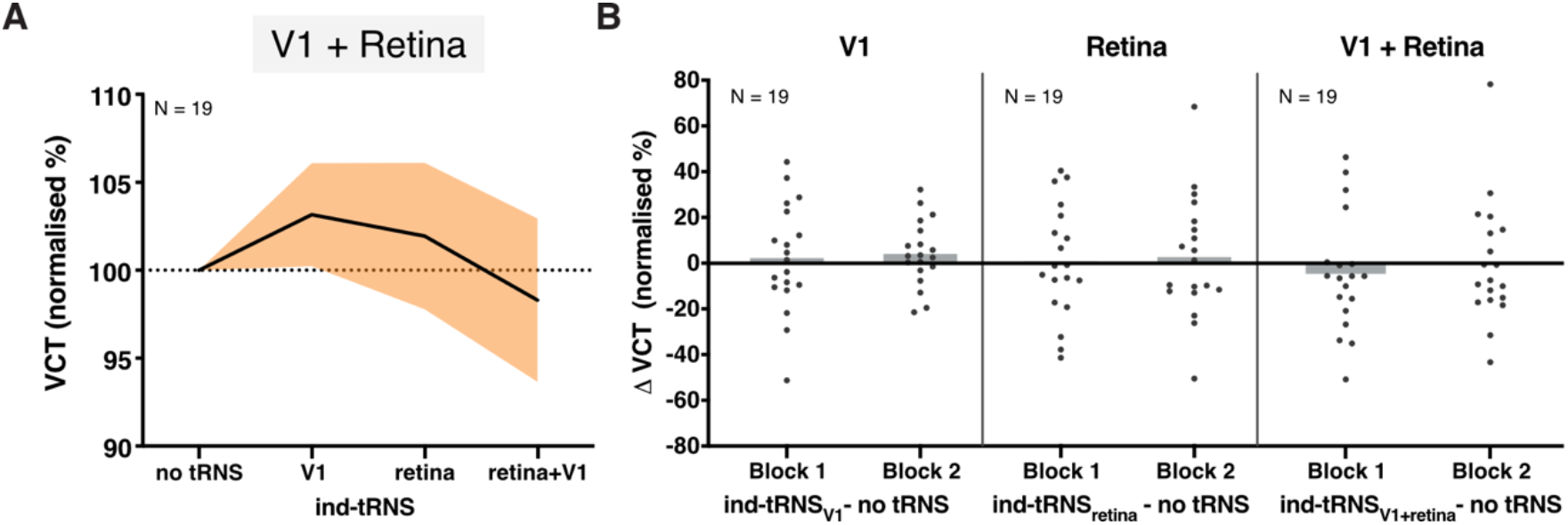
Results of V1+Retina session. **A.** Effect of individualized tRNS_V1_, tRNS_retina_, and tRNS_V1+retina_ on VCT on a group level measured across blocks 1 and 2 in V1+Retina session. All data mean ± SE. **B.** The detection modulation during participants’ optimal ind-tRNS_V1_, ind-tRNS_retina_, and simultaneous ind-tRNS_V1+retina_ in blocks 1 and 2 of V1+Retina session. Gray dots indicate single subject data, gray bar indicates group mean.

In the 3^rd^ session we also retested the effects of individually optimized tRNS intensities defined in V1 and Retina sessions. The effect of ind-tRNS_V1_ found in V1 session was not reproduced between sessions when VCT was measured during ind-tRNS_V1_ in session 3 (t_(18)_ = −0.18, p = 0.43, VCT decrease in 9 out of 19 individuals, MD = 2.24 ±23.63%, and t_(18)_ = −1.37, p = 0.09, VCT decrease in 6 out of 19 individuals, MD = 4.1 ±14.28%, in the 1^st^ and 2^nd^ block respectively, **Figure 6B**). There was also no association between behavioral improvements measured during ind-tRNS_V1_ in the 1^st^ blocks of V1 and V1+Retina sessions (r = 0.12, p = 0.961, N=19), indicating that once-optimized tRNS intensity does not lead to consistent effects between sessions. Similarly to Retina session, participants’ ind-tRNS_retina_ did not lower the VCT compared to the no tRNS condition when retested on the VCT data in session 3 (t_(18)_ = 0.12, p = 0.45, VCT decrease in 11 out of 19 individuals, MD = 1.02% ± 24.57%, and t_(18)_ = - 0.17, p = 0.44, VCT decrease in 9 out of 19 individuals, MD = 2.91 ±26.51%, in the 1^st^ and 2^nd^ block respectively, **Figure 6B**). There was also no association between behavioral improvements measured during ind-tRNS_retina_ in the 1^st^ blocks of Retina and V1+Retina sessions (r = −0.252, p = 0.297, N=19).

## 4. Discussion

The present study investigated the effects of tRNS on visual contrast sensitivity, when applied to different neuronal substrates along the retino-cortical pathway. We measured VCT during tRNS_V1_ and tRNS_retina_ and tRNS_V1+retina_ across 3 experimental sessions. We found consistent tRNS-induced enhancement of visual contrast detection during V1 stimulation (**Figure 5A-C**). but not retina stimulation (**Figure 5D-F**). We also did not observe any additive effects on contrast detection when noise stimulation was simultaneously applied to V1 and retina (**Figure 6A, B**). The online modulation effects of individually optimized tRNS_V1_ intensities were replicated within session (i.e., across two separate blocks) (**Figure 5C**), but not between experimental sessions (**Figure 6B**). Our findings likely reflect acute effects on contrast processing rather than after-effects, as stimulation was only applied for short intervals (2 s) and always interleaved with control (no tRNS) conditions.

### 4.1 tRNS improves visual sensitivity in V1

Our findings confirm previous evidence that the detection of visual stimuli is enhanced when tRNS is added centrally to V1 at optimal intensity (**Figure 5A**; van der Groen and Wenderoth, 2016) even though a different outcome measurement was used (i.e., VCT instead of detection accuracy of subthreshold stimuli). As such, it constitutes to a conceptual replication of the earlier study. The modulation observed here was characterized by large effect size (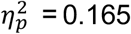, **Figure 5A**), stronger than the intermediate effect size (Cohen’s d = 0.77) found by van der Groen and Wenderoth (2016). Thus, the threshold tracking procedure (Brainard, 1997; Kleiner et al., 2007; Pelli, 1997; Watson and Pelli, 1983) used in our experiments seems to provide a sensitive and reliable estimate of behavioral effects of tRNS_V1_. Moreover, the 4-AFC task protocol used in our study was shown to be more efficient for threshold estimation than commonly used 2-AFC (Jäkel and Wichmann, 2006).

It has been argued previously that tRNS benefits visual detection via the stochastic resonance (SR) mechanism, i.e. the detection probability of weak, subthreshold signals in nonlinear systems can be enhanced if optimally adjusted random noise is added (McDonnell and Abbott, 2009; Moss et al., 2004). One important feature indicative of the SR phenomenon is that noise benefits are a function of noise intensity and exhibit an inverted U-shape relationship, i.e. while the optimal level of noise benefits performance, excessive noise is detrimental (Pavan et al., 2019; van der Groen et al., 2018; van der Groen and Wenderoth, 2016). In the V1 session, we could show that task performance accuracy changed according to an inverted-U-shaped function with increasing tRNS_V1_ intensities (ranging from 0 to 1.5mA, **Figure 5A**) which is consistent with a SR mechanism. Visual detection enhancement was reflected in lower VCT and this improvement was most likely driven by effective stimulation of V1 rather than unspecific tRNS effects such as cutaneous stimulation and associated effects on arousal, as confirmed in the additional analysis using cutaneous sensation detection during tRNS as a covariate.

Based on the behavioral task performance, we determined which tRNS_V1_ intensity was optimal on the individual level (i.e., ind-tRNS_V1_ causing the lowest VCT for each participant, **Figure 5B**). The optimal noise intensities varied across individuals, similar to effects previously shown for noise added both to the stimulus (Collins et al., 1996; Martínez et al., 2007) or centrally to V1 (van der Groen and Wenderoth, 2016). Even though the ind-tRNS_V1_ intensities defined separately in 1^st^ and 2^nd^ blocks of V1 session were not correlated, we demonstrated that the ind-tRNS_V1_ (from 1^st^ block) results in consistent online enhancement effects when retested on the independent data set (VCT in 2^nd^ block) within the experimental session (**Figure 5C**). This indicates that an individually optimized tRNS_V1_ intensity can be considered stable and effective when applied across multiple blocks of a measurement. Notably, the effect of ind-tRNS_V1_ was not replicable on different session (see *Intersession variability in the effects of individualized tRNS protocol on contrast sensitivity* below).

Our study contributes to the evidence for SR as a mechanism underlying online visual processing modulation when tRNS is applied to neural networks in human cortex (Battaglini et al., 2020, 2019; Pavan et al., 2019; van der Groen et al., 2019, 2018; van der Groen and Wenderoth, 2016).

### 4.2 Inconsistencies in the effects of noise on retinal processing of contrast

The present study did not demonstrate systematic noise benefits at the level of the retina. Thus, suggesting that previously reported SR effects on contrast detection might derive mainly from cortical rather than retinal processing. It also shows that SR effects might differ based on the specific characteristic of the stimulated neural tissue.

In our study, we targeted the retina bilaterally with tRNS, to investigate its effects on contrast sensitivity. Although increases in tRNS_retina_ intensity resulted in decreases in VCT, reflecting relative task performance improvements (**Figure 5D**), the effects did not reach statistical significance. Similar to tRNS_V1_, the effects of tRNS_retina_ were variable across study participants. However, even individually determined optimal intensities of tRNS_retina_ did not result in consistent visual processing improvements when retested in separate blocks, both within or between sessions (**Figure 5F**).

Why did tRNS improve contrast detection when applied to V1 but not when applied to retina? In contrast to V1, the retina is characterized by much larger temporal frequency bandwidth toward which it is responsive. One study measured cat ganglion cell responsivity towards temporal frequencies ranging from 0.1 to 100Hz (Frishman et al., 1987). Further studies have shown a similar range of temporal frequency bandwidth in monkey retina (Benardete and Kaplan, 1999) and even higher cut-off frequencies in response to electrical stimulation in rabbit retina (Cai et al., 2011). Moreover, a fMRI study in humans showed a much higher temporal frequency bandwidth cut-off in human Lateral Geniculate Nucleus (LGN – recipient of retinal ganglion cells’ signals) compared to human V1 (Bayram et al., 2016), where the strongest effects are observed for narrow bandwidth of around 4-8Hz (Fawcett et al., 2004). Taken together, stimulus processing at the level of the retina seems to cover a much wider range of temporal frequencies than in V1 and to be more variable. Thus, it is possible that the range of tRNS frequencies used in our experiments, i.e., 100-640 Hz might have been too close to the intrinsic signaling frequencies in the retinal circuitry and in ganglion cells to induce the typical SR effect. V1 neurons, by contrast, respond to frequencies which are one to two magnitudes lower than the tRNS frequencies, and therefore, larger noise benefits could be observed.

Alternatively, the weak effects of tRNS_retina_ might simply be due to filtering properties of retinal neurons. A recent study utilized amplitude modulated tACS (AM-tACS) applied to the retina to investigate the efficacy of different carrier frequencies to induce phosphenes. AM-tACS waveforms comprised of different carrier (50Hz, 200Hz, 1000Hz) and modulation frequencies (8Hz, 16Hz, 28Hz). They found that from the conditions using different carrier frequencies only the lowest one was able to induce phosphenes (Thiele et al., 2021). Thus, suggesting the low-pass nature of retinal neurons which greatly reduces the stimulation effectiveness of evoking suprathreshold response (Deans et al., 2007; Thiele et al., 2021). The researchers point out, however, that their findings do not rule out potential sub-threshold modulations of neural activity during AM-tACS with high carrier frequencies.

In the Retina session we observed gradual decrease of VCT with increasing tRNS_retina_ intensity on the group level (**Figure 5D**). Even though this effect was not significant, we cannot exclude that VCT would decrease further when higher tRNS_retina_ intensities were used. We have based our stimulation intensities on studies utilizing repetitive transorbital alternating current stimulation with similar intensities (Fedorov et al., 2011; Gall et al., 2011, 2010) and demonstrated improved vision in patients with damaged optic nerve (see also Sabel et al., 2020). However, it is possible that the induced current is more strongly attenuated in our study (which used much higher stimulation frequencies) due to the filter properties of retinal neurons. Moreover, in these studies the alternating current was delivered using set of four electrodes positioned above and below participants’ eyes. Such electrodes placement results in different direction of the current and related orientation of the induced electric field than bilateral placement used in this study (**Figure 2A**). Thus, electrodes montage used here might have been suboptimal for retinal ganglion cell stimulation (Dmochowski et al., 2012; Lee et al., 2021).

In summary, we found no evidence that tRNS affects contrast detection at the retinal level. This is interesting from a methodological perspective since it rules out that applying tRNS over V1 elicits confounding effects in the retina, as previously discussed for tACS experiments (Schutter, 2016; Schutter and Hortensius, 2010).

### 4.3 Intersession variability in the effects of individualized tRNS protocol on contrast sensitivity

The influence of individually optimized tRNS on VCT, defined separately for both V1 and the retina in experimental sessions 1 and 2, were retested in session 3. The effects of neither ind-tRNS_V1_, nor ind-tRNS_retina_ were replicated (**Figure 7A-B**). This indicates that optimal tRNS intensity for maximum task performance improvement needs to be individually re-adjusted on each experimental session. These results confirm the well-known variability in the effectiveness of non-invasive brain stimulation (Polanía et al., 2018) and the necessity of carefully designing optimal protocols (Bergmann et al., 2016; Bergmann and Hartwigsen, 2020). The differences in effectiveness of preselected tRNS intensities could result from intrinsic factors such as the participants’ arousal levels or attentional states. Additionally, even though we made sure that our procedure was well standardized, there might have been slight differences in the precise electrodes montage or amount of electroconductive gel, potentially resulting in variability of the electric field induced by tRNS of selected intensity across sessions (Polanía et al., 2018). It is also worth noting that the substantial delay between V1/Retina sessions, and V1+Retina session (5 months on average) because of the COVID-19 pandemic (Bikson et al., 2020) could have also influenced this variability. As the modulation of VCT with ind-tRNS_V1_ or ind-tRNS_retina_ was not replicated in session 3, we also did not observe consistent beneficial additive effects of ind-tRNS delivered simultaneously to V1 and the retina (**Figure 8A-B**).

### 4.4 Conclusions

Our study confirms previous findings that tRNS might enhance visual signal processing of cortical networks via the SR mechanism (Potok et al., 2021; van der Groen and Wenderoth, 2016). When probing the effects of tRNS on contrast sensitivity along the retino-cortical pathway, we demonstrated that V1 seems to be more sensitive than the retina to tRNS-induced modulation of visual processing. Moreover, we found that the individual optimal tRNS intensity applied to V1 to enhance contract detection appears to vary across sessions. The appropriate adjustment of optimal tRNS intensity is therefore important to consider when designing tRNS protocols for perceptual enhancement.

## Acknowledgements

We thank Gabrielle Zbären for her valuable help and feedback on the manuscript and all the participants for their time and effort.

